# *SMN1* copy-number and sequence variant analysis from next generation sequencing data

**DOI:** 10.1101/2020.03.31.014589

**Authors:** Daniel Lopez-Lopez, Carlos Loucera, Rosario Carmona, Virginia Aquino, Josefa Salgado, Sara Pasalodos, María Miranda, Ángel Alonso, Joaquín Dopazo

**Affiliations:** Clinical Bioinformatics Area. Fundación Progreso y Salud (FPS). CDCA, Hospital Virgen del Rocio. 41013. Sevilla. Spain; Computational Systems Medicine, Institute of Biomedicine of Seville (IBIS), Hospital Virgen del Rocio. 41013. Sevilla. Spain; Genomic Medicine Unit. Navarrabiomed. 31008 Pamplona, Navarra. Spain; Department of Genetics. Complejo Hospitalario de Navarra. 31008 Pamplona, Navarra. Spain; Bioinformatics in Rare Diseases (BiER). Centro de Investigación Biomédica en Red de Enfermedades Raras (CIBERER), FPS, Hospital Virgen del Rocío. 41013. Sevilla, Spain; FPS/ELIXIR-es, Hospital Virgen del Rocío, Sevilla, 42013, Spain

## Abstract

Spinal Muscular Atrophy (SMA) is a severe neuromuscular autosomal recessive disorder affecting 1/10,000 live births. Most SMA patients present homozygous deletion of *SMN1*, while the vast majority of SMA carriers present only a single *SMN1* copy. The sequence similarity between *SMN1* and *SMN2*, and the complexity of the SMN locus makes the estimation of the SMN1 copy-number by next generation sequencing (NGS) very difficult Here, we present SMAca, the first python tool to detect SMA carriers and estimate the absolute SMN1 copy-number using NGS data. Moreover, SMAca takes advantage of the knowledge of certain variants specific to *SMN1* duplication to also identify silent carriers. This tool has been validated with a cohort of 326 samples from the Navarra 1000 Genomes project (NAGEN1000). SMAca was developed with a focus on execution speed and easy installation. This combination makes it especially suitable to be integrated into production NGS pipelines. Source code and documentation are available on Github at: www.github.com/babelomics/SMAca

## Introduction

Spinal Muscular Atrophy (SMA; MIM 253300) is an autosomal recessive disorder caused by degeneration of alpha motor neurons in the anterior horn of the spinal cord, leading to hypotonia, muscular atrophy and weakness of proximal muscles, predominantly affecting the lower extremities (Lunn and Wang, 2008). In most populations SMA is caused by homozygous deletions or, less frequently, mutations or exons 1-6 deletions in the survival motor neuron gene (*SMN1*). The disease severity is determined mainly by a copy gene, *SMN2*. The more *SMN2* copies present, the milder the phenotype usually is. Both genes, located on chromosome 5q13.2, can be distinguished by only five nucleotides (Wirth, et al., 2020).

Most individuals have two copies of each *SMN1* and *SMN2*, however due to the complex genomic structure, gene conversion and rearrangements occur quite frequently in SMN locus leading to copy-number variants (MacDonald, et al., 2014). The majority of SMA patients, have a *SMN1* deletion or gene conversion of *SMN1* into *SMN2*, which results in a homozygous loss of *SMN1* exon 7 or exons 7 and 8. Establishing the *SMN2* copy number is of importance for SMA patients due to the inverse correlation between disease severity and *SMN2* copy number. SMA carriers are symptom-free and can be identified by the presence of only a single *SMN1* exon 7 copy. About 5% of SMA carriers have two SMN1 copies on one chromosome and 0 copies on the other (2+0) being named “silent carriers”. Finally, a variant, referred to as *SMN1/2*Δ*7-8*, contains one or two extra copies of SMN exons 1-6 of *SMN1* or *SMN2*. This variant is often present in individuals with no, or only one, *SMN2*. Although it is frequently found (23%) in Spanish carriers and non-carries, its clinical significance is not yet completely clear (Calucho, et al., 2018). Frequency of SMA carriers in the population is around 1.7-2.1 % (Larson, et al., 2015; Su, et al., 2011), presenting most of them only a single *SMN1* exon 7 copy.

At present, the gold standard genetic testing for SMA is multiplex ligation-dependent probe amplification (MLPA) of *SMN1* and *SMN2* although it can neither identify silent carriers (2+0) nor subtle mutations in *SMN1* (false-negative rate of ∼5%). However, next-generation sequencing (NGS) technology is rapidly becoming a cost-effective approach for clinical testing (Boycott, et al., 2019). Despite the difficulties that the accurate determination of two genes almost identical inherent to short read technologies, some strategies to process NGS data have already been proposed to detect SMA carriers (Feng, et al., 2017; Larson, et al., 2015). However, to our knowledge, there are not freely available tools that could be derive the mutational status of SMA form primary massive sequencing data.

Here, we present a python tool, SMAca, which is the first application that can detect SMA carriers and concomitantly estimate the absolute *SMN1* copy-number form NGS data. Moreover, SMAca can exploit the knowledge variants specific to SMN1 duplication to identify the silent carriers, following the recommendations for SMA carrier testing by the American College of Medical Genetics and Genomics (Prior, et al., 2011).

## Materials and Methods

### NGS data processing

Raw FASTQ files were processed following a standard NGS pipeline. Briefly, after filtering out low quality reads with *fastp* v0.20.0 (Chen, et al., 2018), reads were aligned with *BWA-MEM* v0.7.16a (Li and Durbin, 2009) against the human reference genome GRCh37/h19. Potential PCR duplicates were marked with Picard v2.17.3 (http://broadinstitute.github.io/picard/). Finally, the whole set of BAM files were analyzed with SMAca in a single batch.

### SMN1 copy-number estimation

The availability of a batch of samples allows a more accurate estimation of SMN1 copy-number. SMAca first calculates the raw proportion of *SMN1* reads over the total number of reads covering *SMN1* and *SMN2* at three specific gene positions (denoted as *a, b* and *c*) for each sample (Table 1).

**Table 1.** *SMN1* and SMN2 different nucleotides. Positions in *SMN1* (and the analogous positions in *SMN2*) used to calculate the raw proportion of *SMN1* reads(*D1ij*) over the total number of reads covering *SMN1* and *SMN2* (*D1ij+D2ij*)

| Position | SMN1 | SMN2 |
| --- | --- | --- |
| a | chr5:70247724 | chr5:69372304 |
| b | chr5:70247773 | chr5:69372353 |
| c | chr5:70247921 | chr5:69372501 |

These positions correspond to single nucleotide differences between *SMN1* and *SMN2*. Raw values are then scaled with respect to 20 control genes (Table 2) previously described to have consistent average coverage relative to *SMN1* and *SNM2* (Larson, et al., 2015). Additionally, two genetic variants that have been associated with duplication events in *SMN1* are also screened and reported (Luo, et al., 2014).

**Table 2.**
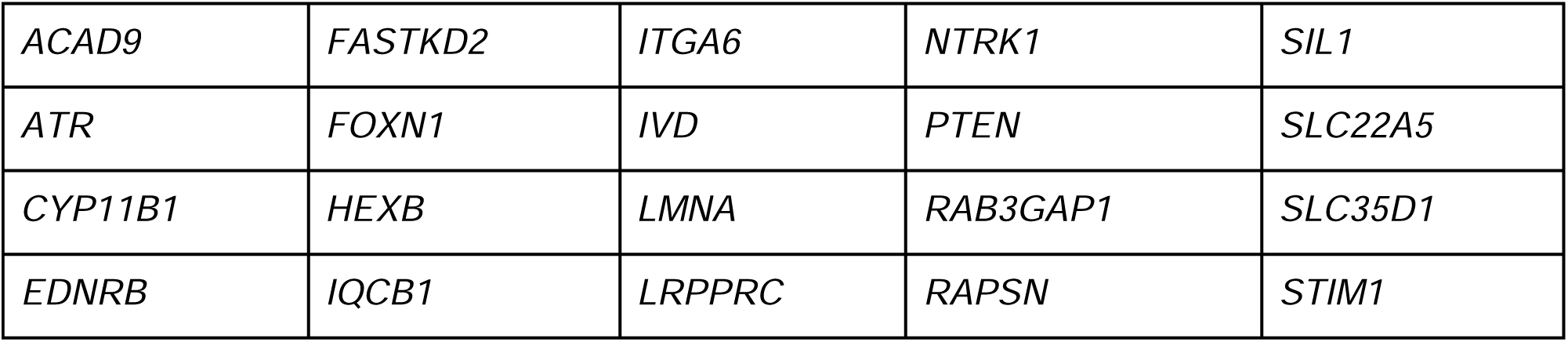
Control genes. List of genes used to calculate the scale factor 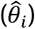 and the scaled proportion of *SMN1* reads (*π*_*ij*_):

In particular, the relative coverage of *SMN1* and *SMN2* with respect to each control gene is calculated: *Z*_*ki*_ *=* (*c*_*i*1_ + *c*_*i*2_)/*H*_*ki*_, were *c*_*i*1_ and c_*i*2_ are the average coverage for the whole genes *SMN1* and *SMN2*, and *H*_*ki*_ is the average coverage for the control gene *k* in the *i*_*th*_ sample. Then, the scale factor 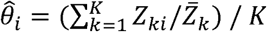, where 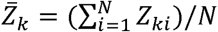, *N* is the total number of samples and *K* the total number of control genes, is calculated for each sample. Finally, raw proportion of *SMN1* reads are scaled: 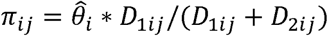, where *D*_1*ij*_ and *D*_2*ij*_ are the raw coverage for *SMN1* and *SMN2* at position j in the *i*_*th*_ sample.

### SMA carrier categorization

Results were classified following some simple rules. Samples with a scaled coverage proportion of SMN1 (*π*_*ij*_) less than ⅓ in positions *a, b* and *c* were marked as likely carriers. The scale factor 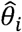(that is proportional to the total *SMN1* and *SMN2* copy-number) and the raw proportion of *SMN1*/*SMN2* depth of coverage at positions *a, b* and *c* (*D*_1*ij*_ /*D*_2*ij*_), where used to estimate the absolute copy-number as follows:

- Genotypes 1 *SMN1*:3 *SMN2* are expected to have 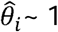 and *D*_1*ij*_ /D_2*ij*_ ∼ ⅓.
- Genotypes 1 *SMN1* :2 *SMN2* are expected to have 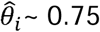 and *D*_1*ij*_ /D_2*ij*_ ∼ ½.
- And genotypes 1 *SMN1*:1 *SMN2* are expected to have 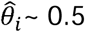 and *D*_1*ij*_ /D_2*ij*_ ∼ 1.

In order to detect silent carriers, we select samples with two polymorphisms (g.27134T>G and g.27706_27707delAT) associated with duplication events in *SMN1* (Luo, et al., 2014). Depending on the number of *SMN2* copies, the expected 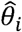 should be close to 0.75 (2:1) or 0.5 (2:0) and, in both cases, the scaled coverage proportion of *SMN1* should be close to ½ in each position.

## Results

### Experimental validation

In order to test the reliability of SMAca predictions in a real scenario we leveraged our participation in the Navarra 1000 Genomes project NAGEN1000 to screen a dataset of 326 genomes. Among them, 7 samples (2.15%) were identified as putative SMA carriers and successfully validated by MLPA (Supplementary File 1). Interestingly, the percentage of the predicted and further confirmed SMA carriers in our dataset fits perfectly to the expected carrier frequency (2.10%) previously described in the bibliography (Su, et al., 2011). Moreover, the genotype estimation for the SMA carrier samples agreed with the experimental validation as shown in Table 3 (except for the case #7 where the genotype could not be estimated). Interestingly, the case #1 corresponds to a SMA carrier with an extra copy of SMN exons 1-6 (*SMN1/2*Δ*7-8*).

**Table 3.** MLPA results. PI_x: scaled proportion of SMN1 reads in position x; cov_x_p: raw coverage of gene x at position p; scale_factor: scale factor 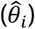 CN_estimation: absolute copy number estimation SMN1:SMN2; MLPA_genotype: genotype inferred from MLPA analysis

| # id | PI_a | PI_b | PI_c | cov SMN1a | Cov SMN1 b_e7 | Cov SMN1 c | Cov SMN2 a | Cov SMN2 b e7 | Cov SMN2 c | Scale factor | CN estimation | MLPA genotype |
| --- | --- | --- | --- | --- | --- | --- | --- | --- | --- | --- | --- | --- |
| 1 | 0,26 | 0,19 | 0,24 | 13 | 8 | 11 | 23 | 23 | 22 | 0,741 | 1:2 | 1:1* |
| 2 | 0,29 | 0,22 | 0,29 | 17 | 14 | 21 | 46 | 54 | 57 | 1,101 | 1:3 | 1:3 |
| 3 | 0,22 | 0,22 | 0,21 | 15 | 15 | 13 | 42 | 41 | 38 | 0,842 | 1:2 | 1:2 |
| 4 | 0,32 | 0,26 | 0,23 | 27 | 18 | 13 | 42 | 39 | 33 | 0,837 | 1:2 | 1:2 |
| 5 | 0,28 | 0,26 | 0,22 | 25 | 24 | 20 | 72 | 78 | 79 | 1,109 | 1:3 | 1:3 |
| 6 | 0,30 | 0,26 | 0,25 | 23 | 22 | 19 | 65 | 74 | 68 | 1,148 | 1:3 | 1:3 |
| 7 | 0,40 | 0,32 | 0,30 | 23 | 19 | 18 | 30 | 36 | 37 | 0,936 | inconclusive | 1:2 |

\* (\*) MLPA analysis showed deletion of exons 7-8 on both genes but three copies of exons 1-6 (impossible to distinguish whether they come from *SMN1* or *SMN2*)

### Performance

With the idea of facilitating the introduction of SMAca in production NGS pipelines it has been optimized for running in different computer environments. A special stress has been made in the parallelization for exploiting multiple cores/processors when available. Figure 1 shows the runtimes with an increasing number of processors. The estimation of SMA mutational and copy number status for 326 genomes from NaGen takes almost one hour and a half in one core but can be reduced to only 3 minutes in 24 cores (see Figure 1)

**Figure 1.**
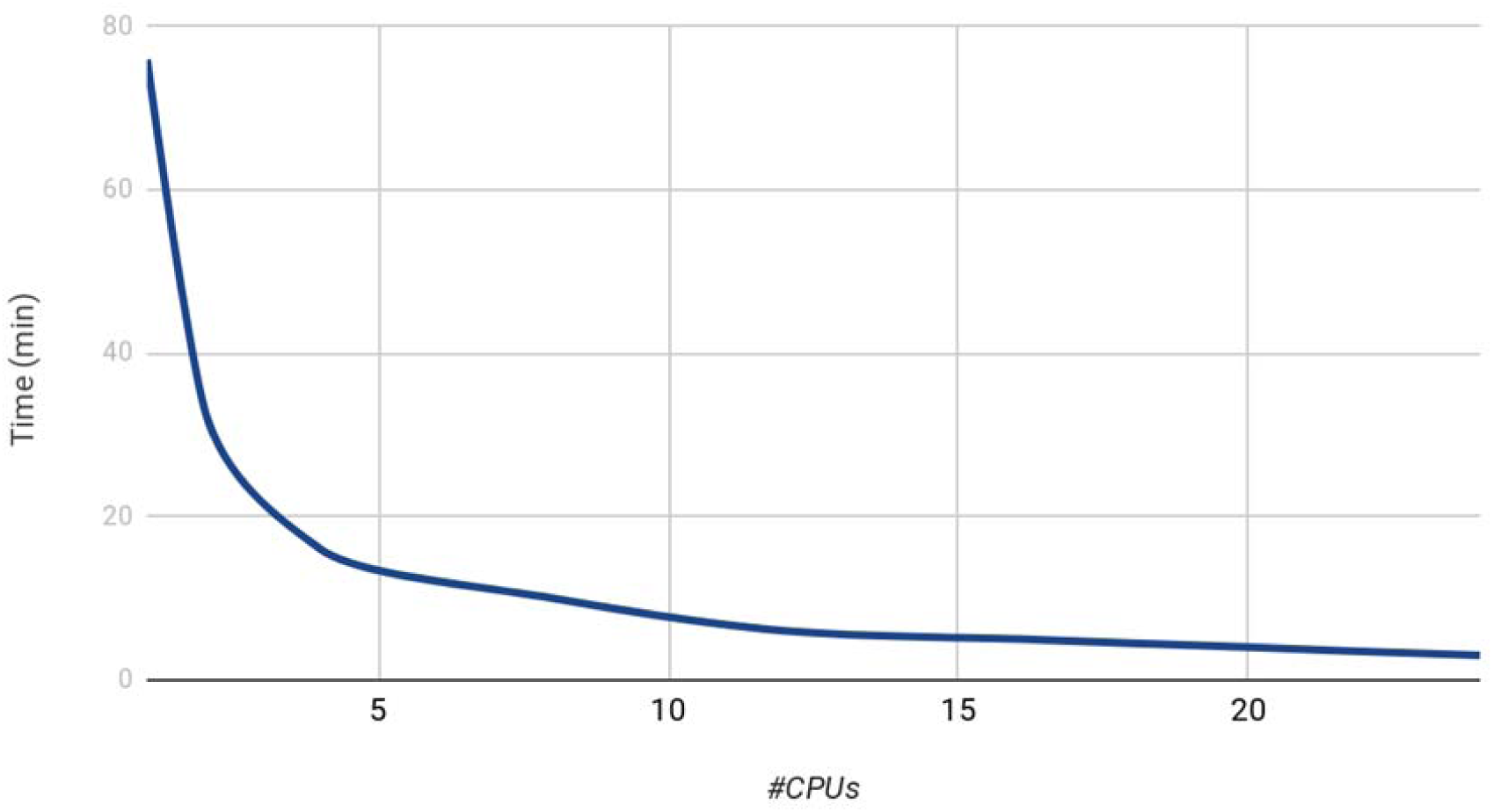
SMAca performance. Elapsed time for the analysis of the dataset (326 whole genome sequences) and different number of CPUs. The analysis of the whole dataset takes only 3 minutes by using 24 threads.

## Conclusions

Here, we present SMAca, the first freely available python tool to detect SMA carriers and estimate the absolute *SMN1* copy-number from NGS data. As a conceptual novelty, SMAca includes the analysis of two polymorphisms that have been linked to silent carriers (Luo, M. et al, 2014) and are recommended for SMA carrier testing by the American College of Medical Genetics and Genomics (Prior, et al., 2011). This tool was developed with a focus on execution speed and easy installation. This combination makes of SMAca an especially attractive tool to be integrated into production NGS pipelines.

## Acknowledgements

This work is supported by grants SAF2017-88908-R from the Spanish Ministry of Economy and Competitiveness and “Plataforma de RecursosBiomoleculares y Bioinformáticos” PT17/0009/0006 from the ISCIII, both co-funded with European Regional Development Funds (ERDF) as well as H2020 Programme of the European Union grants Marie Curie Innovative Training Network “Machine Learning Frontiers in Precision Medicine” (MLFPM) (GA 813533) and “ELIXIR-EXCELERATE fast-track ELIXIR implementation and drive early user exploitation across the life sciences” (GA 676559).

