## Supplementary Figure 1 for "*SMN1* copy-number and sequence variant analysis from next generation sequencing data"

### Supplementary material

#### Materials and Methods

##### Multiplex Ligation-dependent Probe Amplification (MLPA) analysis

The SALSA MLPA probemix P021-B1 SMA-v02 kit is a multiplex PCR technique that uses a single primer pair to amplify up to 32 probes, each with a unique genomic target and length between 175 and 445nt. Four probes are specific for sequences in exon 7 or 8 of either *SMN1* or *SMN2*. 17 probes detect sequences present in both *SMN1* and *SMN2*. There is one probe for the *NAIP* gene and ten reference probes. PCR amplicons are fluorescently labelled and separated and quantified by capillary electrophoresis (Applied Biosystems 3500 Genetic Analyzer) following the MLPA® General Protocol (MRC-Holland). By comparing the resulting peak pattern of a sample to those of a set of reference samples, the number of genomic targets present in the sample of interest can be determined (Coffalyser MLPA analysis software; MRC-Holland). Coffalyser displays lower border (red) and upper border (blue) in ratio charts. In general, when a probe ratio crosses these borders, it is indicative for a duplication or deletion, assuming that the normal copy number of the sequence targeted by the probe is two (Coffalyser Reference Manual, MRC-Holland).

**MLPA results for samples identified as putative SMA carriers**

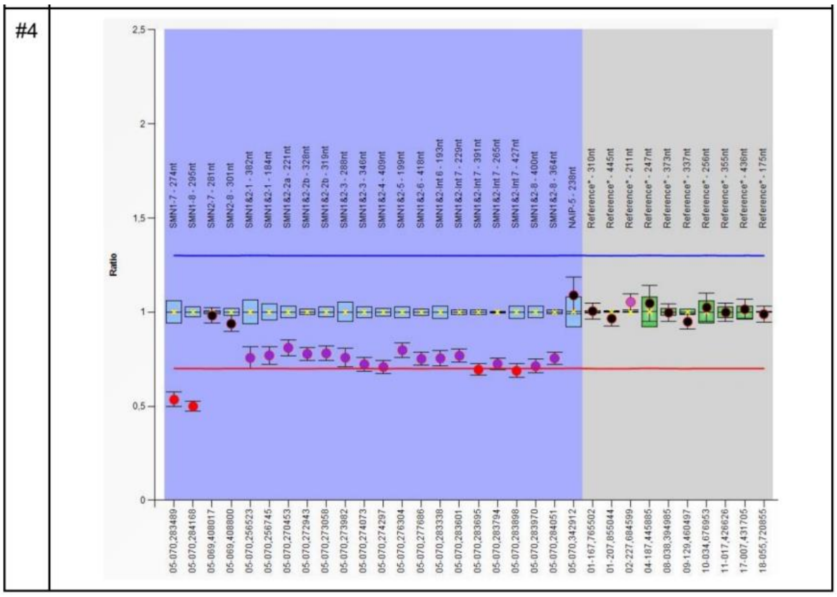


**Figure 1:** SMA carrier with a single *SMN1* copy and two copies of *SMN2* (cases #7 and #4)

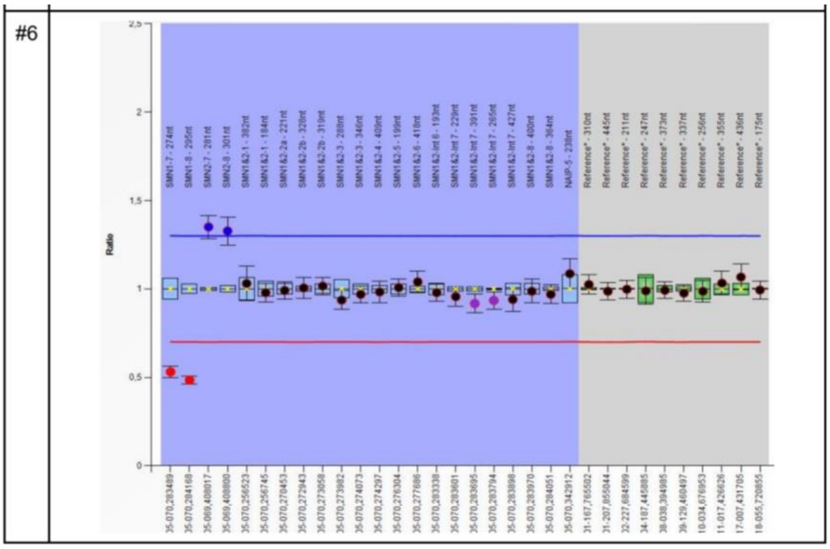


**Figure 2:** SMA carrier with a single *SMN1* copy and three copies of *SMN2* (cases #2, #5 and #6)

| id | MLPA result |
|---|---|
| #1 | 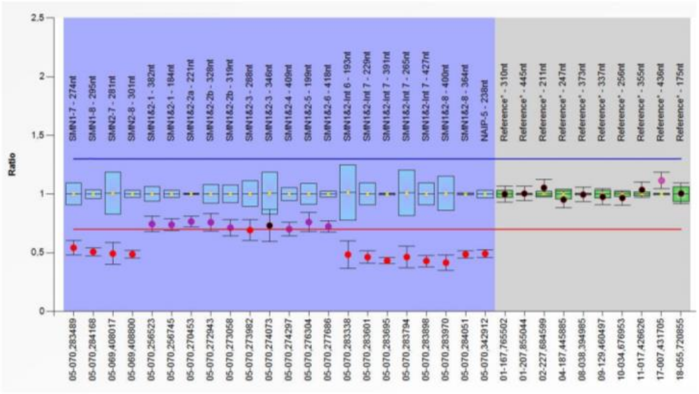 |

**Figure 3:** SMA carrier with a single *SMN1* copy, a single copy of *SMN2* and an extra copy of exons 1-6 of *SMN1* or *SMN2* (*SMN1/2Δ7-8)* (case #1)

| | |
|---|---|
| #3 | 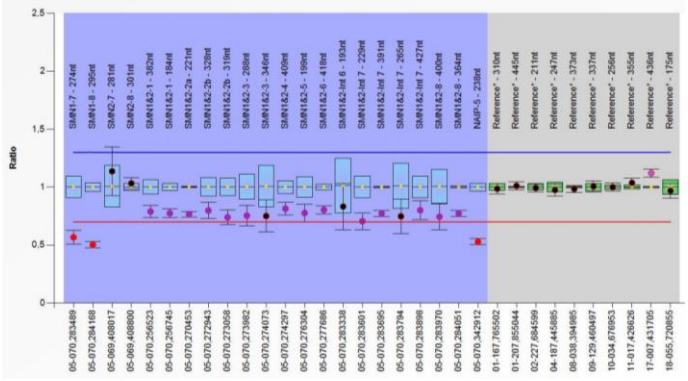 |

**Figure 4:** SMA carrier with a single *SMN1* copy, two copies of *SMN2* and one copy of NAIP gene (exon 5) (case #3). Some publications indicate that patients with fewer copies of NAIP have more severe phenotypes than patients with more copies.
